# Mesophyll specific expression of a bacterial mercury transporter-based vacuolar sequestration machinery sufficiently enhances mercury tolerance of Arabidopsis

**DOI:** 10.1101/2022.07.04.498776

**Authors:** Shimpei Uraguchi, Yuka Ohshiro, Mayuu Okuda, Shiho Kawakami, Nene Yoneyama, Yuta Tsuchiya, Ryosuke Nakamura, Yasukazu Takanezawa, Masako Kiyono

## Abstract

We aimed to efficiently enhance plant Hg(II) tolerance by the transgenic approach utilizing a bacterial mercury transporter MerC, an Arabidopsis mesophyll specific promoter *pRBCS1A*, and a vacuolar membrane targeting syntaxin AtVAM3/SYP22. We generated two independent homozygous Arabidopsis pRBCS1A-TCV lines expressing *mT-Sapphire-MerC-AtVAM3* under the control of *pRBCS1A*. Quantitative RT-PCR showed that the transgene was expressed specifically in shoots of pRBCS1A-TCV lines. Confocal analyses further demonstrated the leaf mesophyll specific expression of mT-Sapphire-MerC-AtVAM3. Confocal observation of the protoplast derived from the F1 plants of the pRBCS1A-TCV line and the tonoplast marker line p35S-GFP-δTIP showed the tonoplast colocalization of mT-Sapphire-MerC-AtVAM3 and GFP-δTIP. These results clearly demonstrated that mT-Sapphire-MerC-AtVAM3 expression in Arabidopsis is spatially regulated as designed at the transcript and the membrane trafficking levels. We then examined the Hg(II) tolerance of the pRBCS1A-TCV lines as well as the p35S-driven MerC-AtVAM3 expressing line p35S-CV under the various Hg(II) stress conditions. Short-term (12 d) Hg(II) treatment indicated the enhanced Hg(II) tolerance of both pRBCS1A-TCV and p35S-CV lines. The longer (3 weeks) Hg(II) treatment highlighted the better shoot growth of the transgenic plants compared to the wild-type Col-0 and the pRBCS1A-TCV lines were more tolerant to Hg(II) stress than the p35S-CV line. These results suggest that mesophyll-specific expression of MerC-AtVAM3 is sufficient or even better to enhance the Arabidopsis Hg(II) tolerance. The Hg accumulation in roots and shoots did not differ between the wild-type Col-0 and the MerC-AtVAM3 expressing plants, suggesting that the boosted Hg(II) tolerance of the transgenic lines would be attributed to vacuolar Hg-sequestration by the tonoplast-localized MerC. Further perspectives of the MerC-based plant engineering are also discussed.

## INTRODUCTION

Due to its serious toxicity, mercury (Hg) ranks in the top 3 on the US Agency for Toxic Substances and Disease Registry (ATSDR) 2019 Priority List of Hazardous Substances (https://www.atsdr.cdc.gov/SPL/), together with arsenic and lead. Soil contamination by Hg is increasingly reported especially in developing countries, attributable to conventional industrial activities like mining and recent anthropogenic activities such as e-waste recycling (Awasthi et al., 2016). Accordingly, recent studies report contamination of cereal grains and vegetables by the inorganic mercury [Hg(II)] (Li et al., 2017; Tang et al., 2018). It is therefore important to reduce the risk of Hg toxicity to human health, and phytoremediation of Hg-contaminated soils is one of the potential approaches for this purpose.

Superior Hg accumulation and tolerance are the key plant traits to achieve efficient phytoremediation of Hg(II) contaminated environment. However, Hg(II) is not an essential element and plants do not appear to have Hg(II)-specific uptake pathways. Nutritional cation transporters are likely to misrecognize Hg(II) as a non-native substrate and mediate Hg(II) uptake by roots. Therefore, molecular manipulation of plant endogenous transporters does not appear to be a potential approach to enhance plant Hg uptake and accumulation for phytoremediation. To overcome this problem, we have been focusing on a bacterial mercury transporter MerC. MerC gene is identified among *mer* operons that confer mercury resistance to bacteria (Kusano et al., 1990; Sahlman et al., 1997; Inoue et al., 2014). Although there are three other Mer transporters (MerE, MerF, and MerT), we found the highest inward mercury transport activity for MerC (Sone et al., 2013). These results suggest MerC as a potential molecular tool to enhance plant Hg(II) uptake and accumulation by its ectopic expression in plants, however, GFP-MerC is mainly stuck at ER-Golgi in Arabidopsis cells and not localized to plasma-membrane (Kiyono et al., 2012). But when fused with SYP121 which is a plasma-membrane resident Arabidopsis syntaxin, MerC is mainly localized to plasma-membrane in Arabidopsis (Kiyono et al., 2012, 2013). p35S-driven ubiquitous expression of MerC-SYP121 increased Hg accumulation in Arabidopsis, probably attributed to enhanced Hg uptake in roots through plasma-membrane localized MerC (Kiyono et al., 2013). Moreover, our recent studies demonstrate that root surface or root endodermis specific expression of MerC-SYP121 sufficiently enhances Hg accumulation in Arabidopsis shoots, as like the case for the p35S-driven ubiquitous MerC-SYP121 expression (Uraguchi et al., 2019a, 2019b).

These successes of the MerC-SYP121 expression system imply another possibility that MerC can also function in plant vacuolar sequestration of Hg for its detoxification. To test this hypothesis, another syntaxin AtVAM3/SYP22 was used to regulate MerC subcellular localization to the tonoplast. AtVAM3 is a tonoplast-resident syntaxin (Sato et al., 1997), and MerC-AtVAM3 fusion protein is localized to the tonoplast in Arabidopsis (Kiyono et al., 2012, 2013). p35S-driven ubiquitous expression of MerC-AtVAM3 improved Arabidopsis tolerance to mercurial stress (Kiyono et al., 2013). These results suggest that ectopically expressed MerC-AtVAM3 functions as vacuolar Hg-sequestration machinery and mitigates Hg toxicity in plant cells.

For efficient phytoremediation, both higher accumulation of target metals in the shoot and higher tolerance against the metal are desirable plant traits. We are planning to establish a multi-layered MerC expression system to confer both enhanced Hg accumulation and improved Hg tolerance to plants. The root cell-type-specific MerC-SYP121 expression systems pEpi and pSCR can be used as minimum expression tools to boost root Hg uptake (Figure 1A). MerC-SYP121 expression in the root epidermis (pEpi) and in the endodermis (pSCR) efficiently enhances Hg accumulation in Arabidopsis and the boosting effects are comparable to that of p35S-driven system (Uraguchi et al., 2019a, 2019b). This study aimed to examine whether leaf mesophyll specific expression of MerC-AtVAM3, vacuolar Hg-sequestration machinery would sufficiently improve Arabidopsis Hg(II) tolerance (Figure 1A) as previously reported for the p35S-driven ubiquitous expression of MerC-AtVAM3 (Kiyono et al., 2013). We generated Arabidopsis plants expressing mT-Sapphire-MerC-AtVAM3 specifically in the leaf mesophyll cells using the mesophyll specific rubisco-related promoter *pRBCS1A* (Mustroph et al., 2009). We demonstrated that the expression of mT-Sapphire-MerC-AtVAM3 on the tonoplast of leaf mesophyll cells sufficiently enhanced Hg tolerance, and the effect of the transgene was comparable to the previously established p35S-driven MerC-AtVAM3 expressing line. The results suggest that mesophyll-specific expression of a bacterial mercury transporter-based vacuolar sequestration machinery is sufficient to enhance mercury tolerance of Arabidopsis. Further perspectives on pyramiding the cell-type-specific MerC expression systems are discussed.

**Figure 1.**
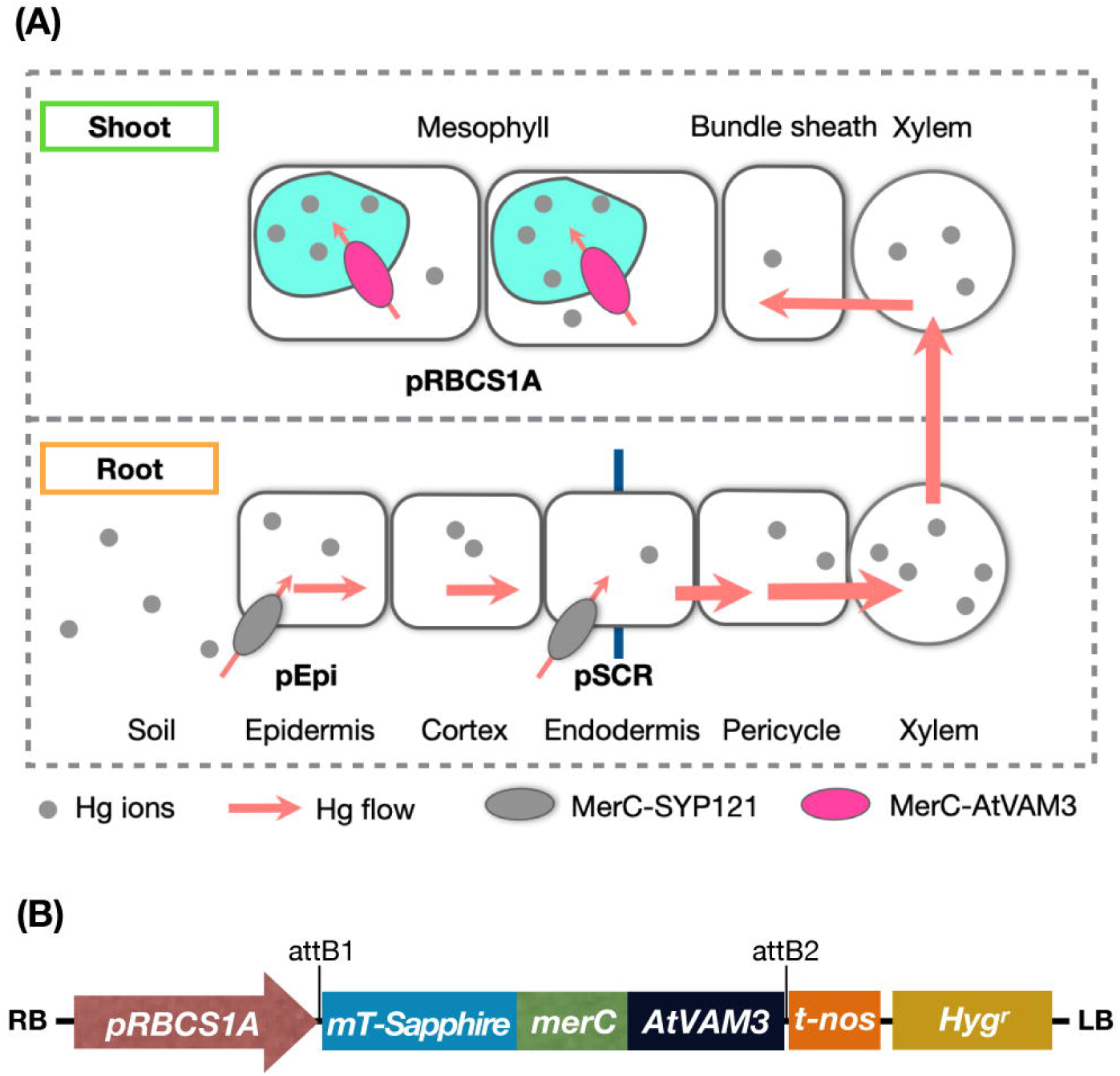
(**A**) Schematic of mercury transport in the bacterial mercury transporter MerC expressing transgenic plants. Root surface-specific (pEpi line) or endodermis-specific (pSCR line) expression of MerC-SYP121 in the plasma-membrane enhances mercury accumulation of the Arabidopsis plants, by boosted Hg(II) uptake in roots (Uraguchi et al., 2019a, 2019b). In shoots, MerC-AtVAM3 in the tonoplast (vacuolar membrane) of the mesophyll cells is expected to mediate vacuolar compartmentation of cytosolic mercury for the detoxification. **(B)** Schematic structure of the T-DNA region in the binary vector constructed for generating pRBCS1A-TCV transgenic plants. RB, right border; *pRBCS1A*, a mesophyll specific promoter; *mT-Sapphire*, a fluorescent reporter gene; *merC*, a bacterial mercury transporter gene; *AtVAM3*, an Arabidopsis tonoplast-resident SNARE gene; t-nos, NOS terminator; Hyg^r^, hygromycin resistance gene; LB, left border.

## MATERIALS AND METHODS

### Plant materials

*Arabidopsis thaliana* Col-0 ecotype was used as wild-type control. For transformation Col-0 was used to obtain *pRBCS1A::mT-Sapphire-merC-AtVAM3* transgenic lines (hereafter “pRBCS1A-TCV lines”) as described below. The ubiquitously-expressing line *p35S::merC*-*AtVAM3* (previously designated as “CV11 line”, hereafter “p35S-CV line” in this study) (Kiyono et al., 2012) was also used.

### Plasmid construction and plant transformation

The primers used for plasmid construction were listed in Supplementary Table S1. A 2-kbp fragment upstream of the start codon of the *RBCS1A* gene (AT1G67090) was amplified from a plasmid harboring the corresponding fragment (Mustroph et al., 2009) which was kindly provided by Prof. Dr. Angelika Mustroph (University of Bayreuth). The obtained DNA fragment, designated as pRBCS1A was ligated into the upstream of the Gateway cassette of pGWB501(Nakagawa et al., 2007) using the SbfI and XbaI sites. The coding sequence of *mT-Sapphire* was amplified from a plasmid mT-Sapphire-C1, a gift from Michael Davidson (Addgene plasmid # 54545). The *merC-AtVAM3* fragment was amplified from pMACV1 (Kiyono et al., 2012). The amplified *mT-Sapphire* and *merC-AtVAM3* fragments shared a 15 bp linker sequence for Gly-Gly-Gly-Gly-Ala in the 3’ and 5’ end of the respective fragments. The fragments were fused by overlap extension PCR and the resultant fragment *mT-Sapphire-merC-AtVAM3* was directionally inserted into pENTR/D-TOPO (Thermo Fisher Scientific). Then *mT-Sapphire-merC-AtVAM3* in pENTR/D-TOPO was subcloned into the pGWB501 harboring *pRBCS1A* upstream of the Gateway cassette using the LR recombination reaction to obtain *pRBCS1A::mT-Sapphire-merC-AtVAM3* (Figure 1B). The resultant plasmid was introduced into *Agrobacterium tumefaciens* GV3101::pMP90, which were then used for the transformation of *Arabidopsis thaliana* Col-0 wild-type plants by the floral dip method (Clough and Bent, 1998).

### Quantitative RT-PCR

To examine the transgene expression, 12-d-old plants were subjected to RNA extraction. NucleoSpin RNA Plant (MACHEREY-NAGEL) was used for total RNA extraction from roots or shoots. DNase treatment was applied during the RNA extraction. PrimeScript RT Master Mix (Takara Bio) was used for cDNA synthesis and quantitative RT-PCR was performed with PowerUp SYBR Green Master Mix (Thermo Fisher Scientific). *AtEF1a* (At5g60390) served as an internal control. *merC* primers were used to detect the transgene transcript. A relative quantification method using standard curves was applied. The primer sequences used for expression analyses are listed in Supplementary Table S1.

### Confocal laser microscopy

The expression of mT-Sapphire-merC-AtVAM3 in roots and leaves of the pRBCS1A-TCV lines was examined by a laser scanning confocal microscope (FV-3000, Olympus). Prior to the observation, roots were stained with 4 µM FM4-64 for 5 min. Chlorophyll autofluorescence was also observed for leaf samples. The excitation and detection wavelengths were 405 nm and 485–535 nm for mT-Sapphire; 488 nm and 600–700 nm for FM4-64 and chlorophyll autofluorescence, respectively.

Subcellular localization of mT-Sapphire-merC-AtVAM3 was further examined in the F1 plants derived from crossing pRBCS1A-TCV line 7 and the tonoplast marker line p35S-GFP-δTIP (Cutler et al., 2000). Protoplast was prepared from mesophyll of the mature F1 plant leaves by the Tape-Arabidopsis Sandwich method (Wu et al., 2009) with a slight modification. The upper epidermal surface of the leaf was carefully affixed to a strip of adhesive Time tape (Hirasawa). The lower epidermal surface of the leaf was then affixed to a strip of less adhesive Magic tape (ASKUL). The Magic tape on the lower surface side of the leaf was then carefully peeled off from the Time tape. This easily removed the lower epidermal surface cell layer, while the rest of the leaf stayed affixed to the Time tape. The peeled leaves (four to five leaves for each preparation) adhering to the Time tape were transferred to a Petri dish containing 10 mL of the enzyme solution [1% cellulase ‘Onozuka’ R10 (Yakult), 0.25% macerozyme ‘Onozuka’ R10 (Yakult), 0.4 M mannitol, 10 mM CaCl_2_, 20 mM KCl, 0.1% BSA, and 20 mM MES, pH 5.7] and gently shaken on a shaker at 70 rpm for 40 min to release the protoplasts into the enzyme solution. To collect the protoplasts, the solution was centrifuged at 100 × *g* for 3 min with a swing rotor, and the cells were washed twice with 25 mL of prechilled modified W5 solution (154 mM NaCl, 125 mM CaCl_2_, 5 mM KCl, 5 mM glucose, and 2 mM MES, pH 5.7). The protoplasts were resuspended in 0.2 mL of modified MMg solution (0.4M mannitol, 15 mM MgCl_2_, and 4 mM MES, pH 5.7) and used for the confocal observation. The excitation and detection wavelengths were 405 nm and 485–535 nm for mT-Sapphire; 488 nm and 500–530 nm for GFP; 488 nm and 600–700 nm for chlorophyll autofluorescence, respectively.

### Hg(II) tolerance assay

One-tenth strength modified Hoagland medium was used for plant cultivation. The basic medium contained 100 μM (NH_4_)_2_HPO_4_, 200 μM MgSO_4_, 280 μM Ca(NO_3_)_2_, 600 μM KNO_3_, 5 μM Fe-N,N’-di-(2-hydroxybenzoyl)-ethylenediamine-N,N’-diacetic acid (HBED), 1% (w/v) sucrose, and `5 mM MES (pH 5.7) (Tennstedt et al., 2009; Kühnlenz et al., 2014). The following essential microelements were also supplemented to the medium (4.63 µM H_3_BO_3_, 32 nM CuSO_4_, 915 nM MnCl_2_, 77 nM ZnSO_4_, 11 nM MoO_3_). Agar Type A (Sigma-Aldrich, catalog no. A4550-500G) was used for medium solidification. We previously demonstrated that this agar reagent is better suited for evaluating Arabidopsis responses to Hg(II) stress compared to the other agar reagents (Uraguchi et al., 2020). The concentration of agar reagents used for medium solidification was 1% (w/v) for the vertical plate assay and 0.6% (w/v) for the horizontal plate assay. HgCl_2_ was added to the medium with different concentrations as Hg(II) treatments (15, 20, and 25 µM for the vertical plate assay and 5 and 20 µM for the horizontal plate assay).

Arabidopsis seeds were surface sterilized and sown on agar plates. After 2 d stratification at 4°C, plants were grown vertically in a long-day-conditioned growth chamber (16 h light/8 h dark, 22°C) for 12 days or horizontally in a short-day-conditioned growth chamber (8 h light/16 h dark, 22°C) for 21 days. For the vertical growth assay, the plants were photographed after the growth period and plant growth was assessed by primary root length and seedling fresh weight measurements. For the horizontal assay, the plants were photographed, and the seedling fresh weight was measured. For higher Hg(II) dose conditions (20 µM), plants developing a pair of true leaves were also counted as Hg(II)-tolerant plants.

### Mercury accumulation analysis

Mercury accumulation in the plants was evaluated under two different Hg(II) conditions. the plants were grown vertically for 10 d under the long-day condition with 1 µM Hg(II) containing one-tenth strength modified Hoagland medium. The longer Hg(II) treatment (3 weeks) was under horizontal and short-day growth conditions using the medium containing 5 µM Hg(II). Roots and shoots of 15 seedlings were separately pooled as a single replicate. At least three replicates were prepared for each experiment and the experiment was independently repeated twice. At harvest, shoot samples were washed with MilliQ water for twice. Root samples were sequentially desorbed for 10 min each in ice-cold MilliQ water, 20 mM CaCl_2_ (twice), 10 mM EDTA (pH 5.7), and MilliQ water (Uraguchi et al., 2017). The harvested roots and shoots were dried at 50°C before acid digestion. Dried plant samples were wet-digested with HNO_3_ and total mercury concentrations in the digested samples were quantified by a cold vapor atomic absorption spectrometer (HG-400, Hiranuma) (Uraguchi et al., 2019b).

### Statistical analyses

The JASP software ver. 0.15 (JASP team, 2022) was used for statistical analyses. The growth data and mercury accumulation data were analyzed by one-way ANOVA, followed by Dunnett’s test (*p* < 0.05) to examine the significance of differences between Col-0 wild-type and the respective transgenic lines.

## RESULTS AND DISCUSSION

For efficient phytoremediation of toxic metals, both uptake in roots and detoxification in shoots are crucial. The former is for metal-extraction efficiency from soils to plants and the latter is for maintaining plant growth under toxic metal stress conditions. One of the successful examples of improving plant mercurial tolerance by a transgenic approach is a series of studies by Meagher’s group. They showed that the ectopic expression of a mercuric ion reductase MerA gene and/or an organomercurial lyase MerB gene isolated from Hg-resistant bacteria conferred Hg tolerance to plants (Bizily et al., 2000, 2003; Ruiz et al., 2003; Hachez et al., 2014). This transgenic approach was very successful, however, it enhances plant Hg tolerance by facilitating the evaporation of the gaseous form of Hg (Hg0) from leaves. Thus, it contains a risk of re-contaminating the surrounding environment and it does not facilitate Hg accumulation. These points are disadvantages when applying the transgenic plants for phytoremediation of Hg contaminated soils.

To seal in the absorbed Hg in the plant tissues, we have developed another transgenic plant approach by utilizing a bacterial influx Hg transporter MerC fused with plant syntaxin proteins SYP121 or AtVAM3/SYP22. In our system, MerC-SYP121 and MerC-AtVAM3 fusion proteins were preferentially localized to the plasma membrane and tonoplast, respectively (Kiyono et al. 2012, 2013). Expression of MerC-SYP121 and MerC-AtVAM3 driven by the 35S-promoter improved Hg accumulation and tolerance of Arabidopsis plants by facilitated Hg uptake by MerC-SYP121 on the plasma membrane and Hg sequestration to vacuoles by MerC-AtVAM3 on the tonoplast, respectively. These phenotypes are desirable for use in phytoremediation of Hg contaminated soils and sealing the absorbed Hg in the plant body, which both contribute to reducing the environmental and health Hg risks. Recently, we further demonstrated that the boosting effect on shoot Hg accumulation by root epidermis and endodermis specific expression of MerC-SYP121 (Figure 1A) (Uraguchi et al., 2019a, 2019b). For improving plant Hg(II) tolerance by the cell-type specific regulation, this study examined whether mesophyll-specific expression of MerC-AtVAM3 on the tonoplast would sufficiently enhance Hg(II) tolerance through facilitating vacuolar Hg sequestration (Figure 1A).

### Generation of pRBCS1A-TCV lines

We prepared a construct to generate Arabidopsis plants expressing MerC-AtVAM3 specifically in leaf mesophyll cells using the mesophyll-specific promoter pRBCS1A (Figure 1B). mT-Sapphire, a monomeric variant of a violet-excitable GFP with long stokes shift (Ai et al., 2008) was fused to the N-terminus of MerC-AtVAM3 as a reporter. The gene expression was controlled under the mesophyll-specific promoter pRBCS1A (Mustroph et al., 2009). We transformed Arabidopsis wild-type Col-0 and obtained 10 independent T1 plants. Among these hygromycin-resistant T1 plants, four lines showed normal growth, and the other lines showed semi-dwarf or dwarf (and sterile) phenotypes (Supplementary Figure S1). We previously found abnormal shoot development phenotypes when introducing *merC-SYP111* to Col-0 (Kiyono et al., 2012), indicating a possibility that expressing *merC* gene fused to plant SNARE genes may cause such growth defects. Nevertheless, as a result of the selection of the pRBCS1A transgenic plants, line 7 and line 3 were established as homozygous T3 lines with standard fertility and hereafter designated as pRBCS1A-TCV lines.

### Mesophyll- and tonoplast-specific expression of mT-Sapphire-MerC-AtVAM3

We then quantified the transgene expression in the pRBCS1A-TCV lines and compared it with the previously established p35S-driven MerC-AtVAM3-expressing line p35S-CV (Kiyono et al., 2012) by quantitative real-time PCR (Figure 2). The *merC* expression in pRBCS1A-TCV line 7 shoot was comparable to that of the p35S-CV line, followed by pRBCS1A-TCV line 3. As expected, roots of pRBCS1A-TCV lines expressed negligible levels of *merC*, whereas p35S-CV roots expressed a substantial level of *merC*. This shoot-specific transgene expression of the pRBCS1A-TCV lines was further confirmed by confocal microscopy (Figure 3A). No visible mT-Sapphire signal was detected in roots, but leaf mesophyll cells showed a positive mT-Sapphire signal. We also examined leaf epidermis, but there was no detectable expression of the mT-Sapphire, suggesting that mT-Sapphire-MerC-AtVAM3 is expressed predominantly in the mesophyll. We further examined the subcellular localization of mT-Sapphire-MerC-AtVAM3, which was expected to be tonoplast-localized, attributed to a tonoplast-resident SNARE molecule AtVAM3 attachment (Kiyono et al., 2013). For this observation, we crossed pRBCS1A-TCV line 7 and a tonoplast marker line p35S-GFP-δTIP (Cutler et al., 2000). Protoplast was prepared from mesophyll of the mature F1 plant by the Tape-Arabidopsis Sandwich method (Wu et al., 2009) and subjected to confocal observation (Figure 3B). GFP-δTIP signal showed clear vacuolar membrane localization, breaking at the points where chloroplasts interfered (indicated by white arrowheads). The mT-Sapphire signal was mostly colocalized with the GFP-δTIP signal, whereas inner vacuolar signal of mT-Sapphire was also slightly observed. The pKa of mT-Sapphire is 4.9 (Ai et al., 2008), suggesting that the fluorescent protein is resistant to acidic conditions like vacuoles which normally have pH 5 to 5.5. Overall, the protoplast observation suggests that mT-Sapphire-MerC-AtVAM3 is partly deposited to vacuoles and probably degraded, but the majority is tonoplast localized in the mesophyll cells. AtVAM3 is a tonoplast-resident syntaxin (Sato et al., 1997). Fusing MerC and AtVAM3 resulted in the tonoplast localization in Arabidopsis cells, demonstrated by the GFP observation of the suspension cells and the western blotting of the p35S-CV plants after sucrose density gradient fractionation (Kiyono et al., 2012, 2013). The presented protoplast observation (Figure 3B) further and clearly demonstrates that AtVAM3-attachment functions in the tonoplast targeting of MerC in Arabidopsis cells.

**Figure 2.**
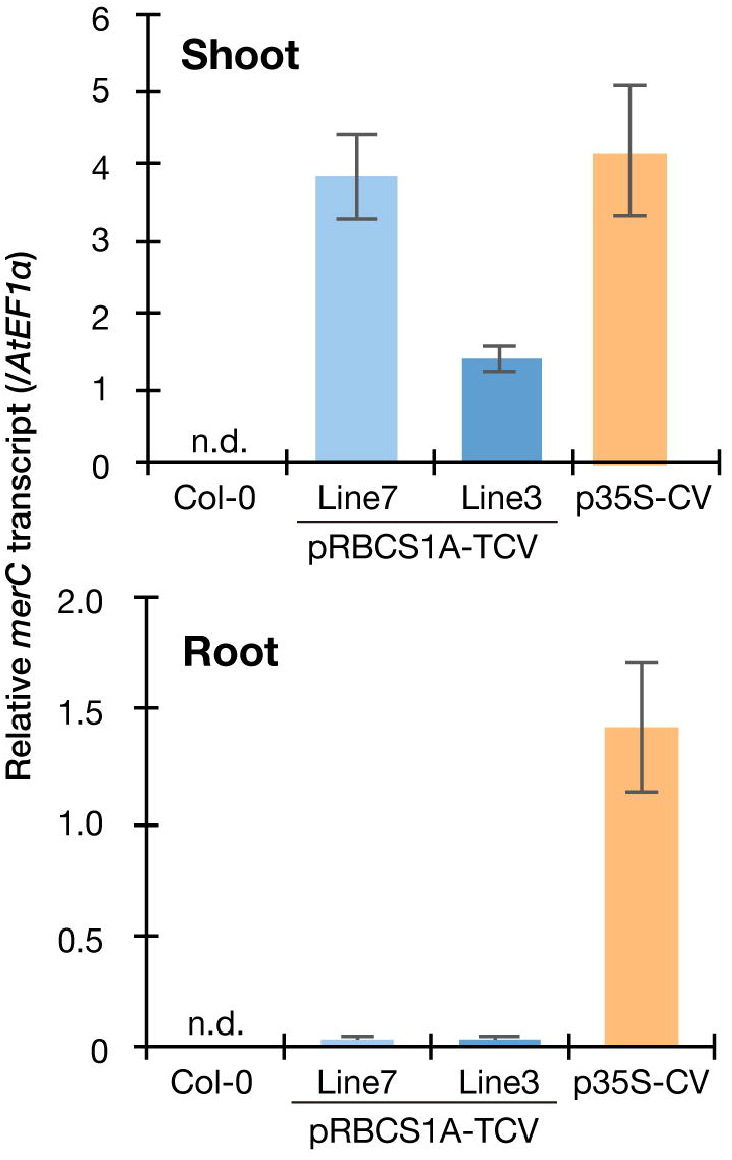
Spatial specificity of the transgene expression in the pRBCS1A-TCV lines by real-time RT-PCR. Expression levels of *merC* were normalized to those of *AtEF1a*. Col-0 wild-type and the previously established ubiquitously expressing line p35S-CV were included for comparison. The data are presented as means with standard errors of four independent biological replicates. n.d., not detected.

**Figure 3.**
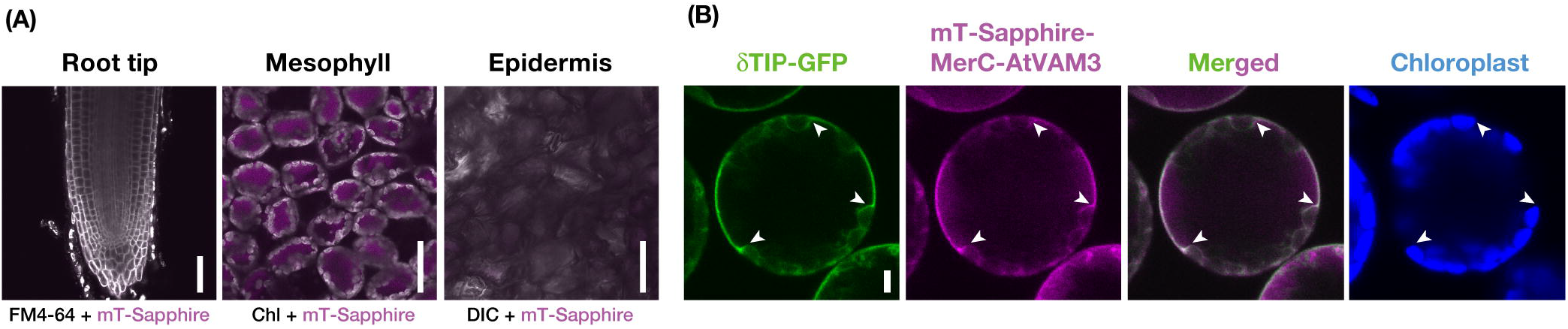
Cell-type specific expression and subcellular localization of mT-Sapphire-MerC-AtVAM3 in the pRBCS1A-TCV plants. **(A)** Mesophyll specific expression of mT-Sapphire-MerC-AtVAM3 in the pRBCS1A-TCV plants. Confocal microscopic images of the root tip, mesophyll, and leaf epidermis of pRBCS1A-TCV line 7 plants were shown. Scale bars = 50 µm. **(B)** Tonoplast localization of mT-Sapphire-MerC-AtVAM3 in the protoplast derived from mesophyll cells of F1 plants of p35S-δTIP-GFP/pRBCS1A-TCV line 7. Scale bar = 5 µm. White arrowheads indicate where the tonoplast signal breaks due to chloroplast interference.

### Hg(II) tolerance of pRBCS1A-TCV lines

We then examined the Hg(II) tolerance of pRBCS1A-TCV lines in comparison to wild-type Col-0 and the ubiquitously expressing line p35S-CV. It was hypothesized that the tonoplast-localized MerC would facilitate vacuolar sequestration of Hg in the cytosol and enhance Hg(II) tolerance of the Arabidopsis plants (Figure 1A). The vertical growth assay demonstrated that the growth of pRBCS1A-TCV lines was comparable to Col-0 under the control agar plate condition but pRBCS1A-TCV lines showed slightly better growth after 15 µM Hg(II) treatment for 12 d (Figure 4A). Fresh weight measurement further showed that root growth of the two pRBCS1A-TCV lines, as well as the p35S-CV line, was significantly better than Col-0 under 15 µM Hg(II) treatment and the tendency was also visible under 20 µM and 25 µM Hg(II) treatments (Figure 4B). However, the difference in the shoot was only significant for pRBCS1A-TCV line 7 under 15 µM Hg(II) treatment in 12-d-old young seedlings. We hypothesized that MerC-AtVAM3 as vacuolar Hg-sequestration machinery would be more functional in mature plants that would develop larger vacuoles.

**Figure 4.**
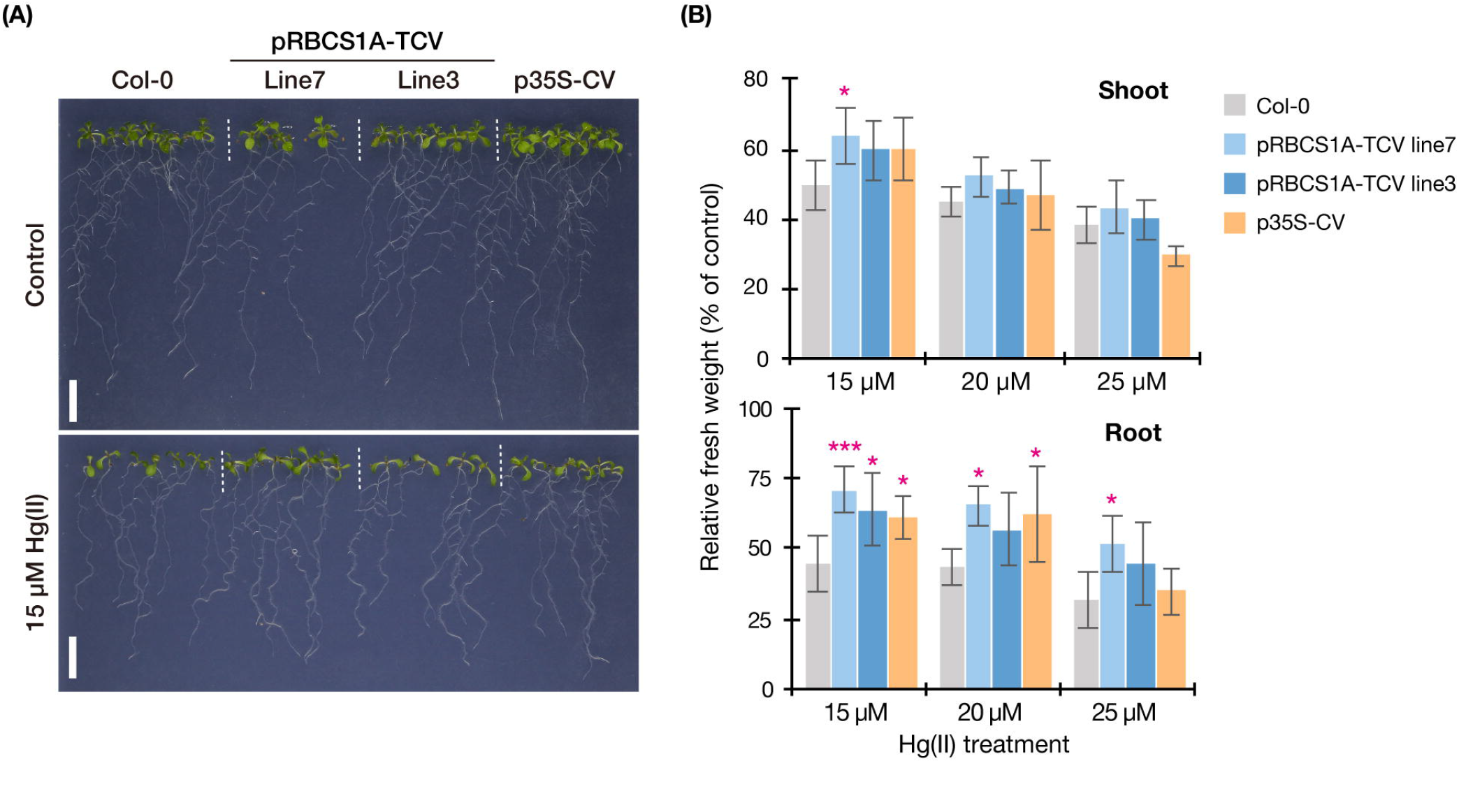
Vertical Hg(II) tolerance assay. Phenotypes **(A)** and seedling growth **(B)** of Col-0, pRBCS1A-TCV lines, and p35S-CV line after different Hg(II) treatments for 12 d. Values are shown as the percentage of each control. Data represent means with SD from two independent experiments with six replicates. Asterisks indicate a significant difference from Col-0 of each treatment (\**P* < 0.05, *** *P* < 0.001, Dunnett’s test).

This possibility was further examined by longer-term horizontal plate assays under a short-day condition which enabled prolonged Hg(II) exposure. After 3 weeks of 5 µM Hg(II) exposure shoot development of Col-0 was inhibited, however, pRBCS1A-TCV line 7 and line 3 were less affected and both showed larger shoot development compared to Col-0 (Figure 5A). The p35S-CV line was similar to Col-0. Shoot fresh weight measurement demonstrated significantly better shoot growth of pRBCS1A-TCV line 7 and line 3 compared to Col-0 under 5 µM Hg(II) treatment, whereas no significant difference was found under the control condition (Figure 5B). We also tested a higher dose of Hg(II) (20 µM), which almost eliminated shoot development of Col-0 (Figure 6A). Under this severe condition, Col-0 rarely developed true leaves and some plants lacked one side of the true leaves pair (indicated with white arrowheads in Figure 6A). On the other hand, the MerC-AtVAM3 expressing lines, including the p35S-CV line, developed a pair of true leaves more frequently than Col-0. We defined a plant with a pair of developed true leaves as a Hg(II) tolerant plant (indicated with asterisks in Figure 6A) and counted the number. Based on the frequency of the Hg(II) tolerant plants, pRBCS1A-TCV line 7 was suggested as the most tolerant line, followed by line 3 and p35S-CV (Figure 6B).

**Figure 5.**
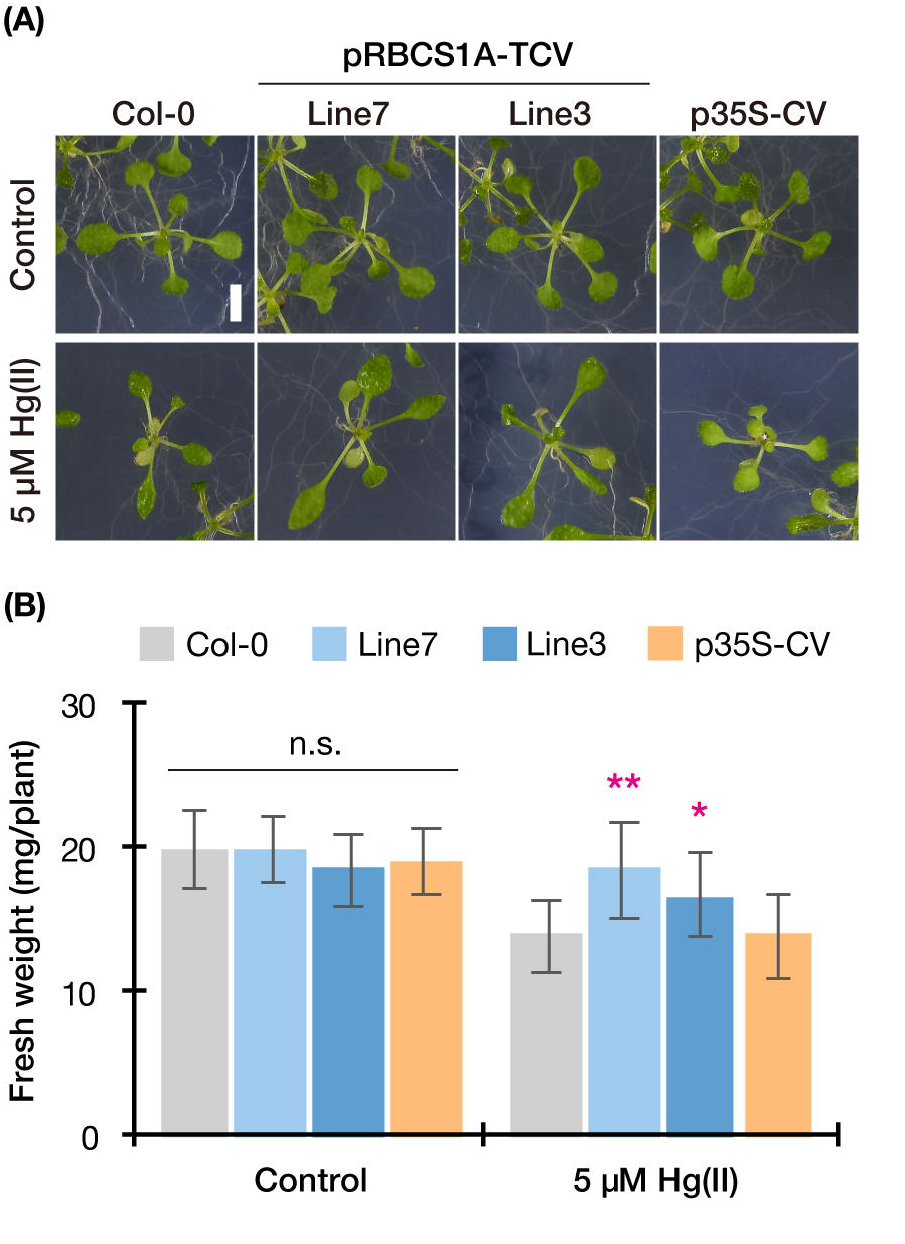
Horizontal Hg(II) tolerance assay with lower dose. Phenotypes **(A)** and shoot growth **(B)** of Col-0, pRBCS1A-TCV lines, and p35S-CV line after 5 µM Hg(II) treatment for 3 weeks. Data represent means with SD from three independent experiments (*n* = 9 – 12). Asterisks indicate a significant difference from Col-0 of each treatment (\**P* < 0.05, ** *P* < 0.01, Dunnett’s test). n.s., not significant.

**Figure 6.**
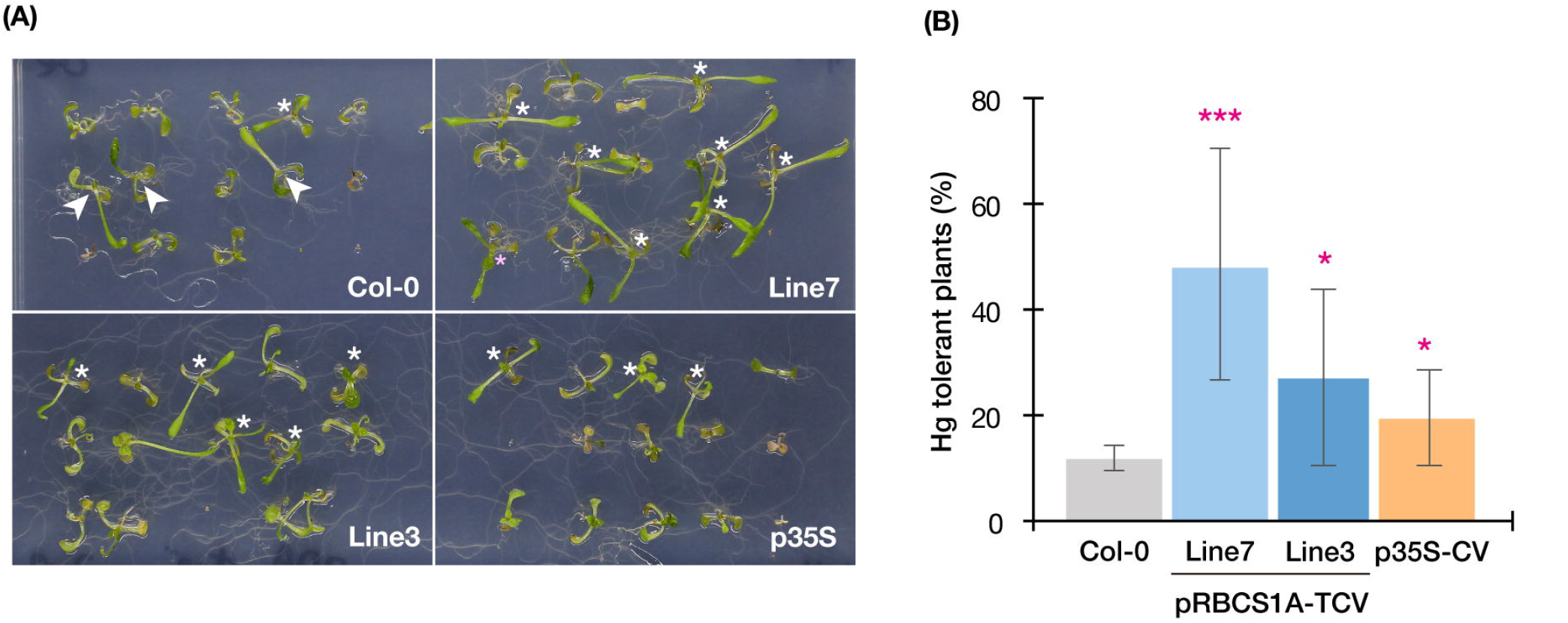
Horizontal Hg(II) tolerance assay with higher dose. **(A)** Phenotypes of Col-0, pRBCS1A-TCV lines, and p35S-CV line after 20 µM Hg(II) treatment for 3 weeks. Asterisks indicate Hg(II)-tolerant plants developing a pair of true leaves. White arrowheads indicate plants lacking one side of the true leaves pair, which were typically observed for Col-0. **(B)** Frequency of Hg-tolerant plants with a pair of true leaves under 20 µM Hg(II) treatment. Data represent means with SD from two independent experiments with eight replicated assays. Asterisks indicate a significant difference from Col-0 (\**P* < 0.05, *** *P* < 0.001, paired t-test).

Taking together with all the Hg(II) tolerance assays (Figures 4-6), the p35S-CV line showed improved Hg(II) tolerance than the Col-0 wild-type as previously reported (Kiyono et al., 2013), however, the pRBCS1A-TCV lines exhibited even better growth under various Hg(II) conditions. This suggests that the tonoplast-targeted expression of MerC-AtVAM3 in mesophyll cells enhances Arabidopsis Hg(II) tolerance more efficiently than the p35S-driven ubiquitous expression system. Leaf mesophyll cells account for a large portion of leaf tissues and would quantitatively function as “sink” of toxic metals taken up by roots and translocated to shoot. It is very likely that the tonoplast-targeted expression of MerC in the mesophyll cells facilitates vacuolar sequestration of Hg and reduces its phytotoxicity (Figure 1A). The importance of the vacuolar sequestration in Hg-detoxification is demonstrated for the Arabidopsis endogenous ABC transporters AtABCC1 and AtABCC2: the loss-of-function mutant *abcc1/abcc2* exhibits the hypersensitivity to mercurials (Park et al., 2012; Uraguchi et al., 2020, 2021). In the present study, we successfully introduced the bacterial mercury transporter to the tonoplast of the mesophyll cells and this vacuolar sequestration machinery was suggested to protect cytosol and other organelles from Hg-toxicity. Given that the enhanced Hg(II)-tolerance of the MerC-AtVAM3 expressing lines is attributed to the subcellular distribution change of mercury from the cytosol to the vacuole, total Hg accumulation in shoots and roots would not differ between Col-0 and the transgenic plants. To examine this hypothesis, we measured Hg accumulation in plants under two different Hg(II) treatments (Figure 7). No obvious difference was observed between Col-0 and the MerC-AtVAM3 expressing lines (Figure 7). This result supports our hypothesis that the improved Hg tolerance of the MerC-AtVAM3 expressing lines is attributed to preferential vacuolar sequestration of Hg, not due to restriction of Hg uptake and/or root-to-shoot accumulation.

**Figure 7.**
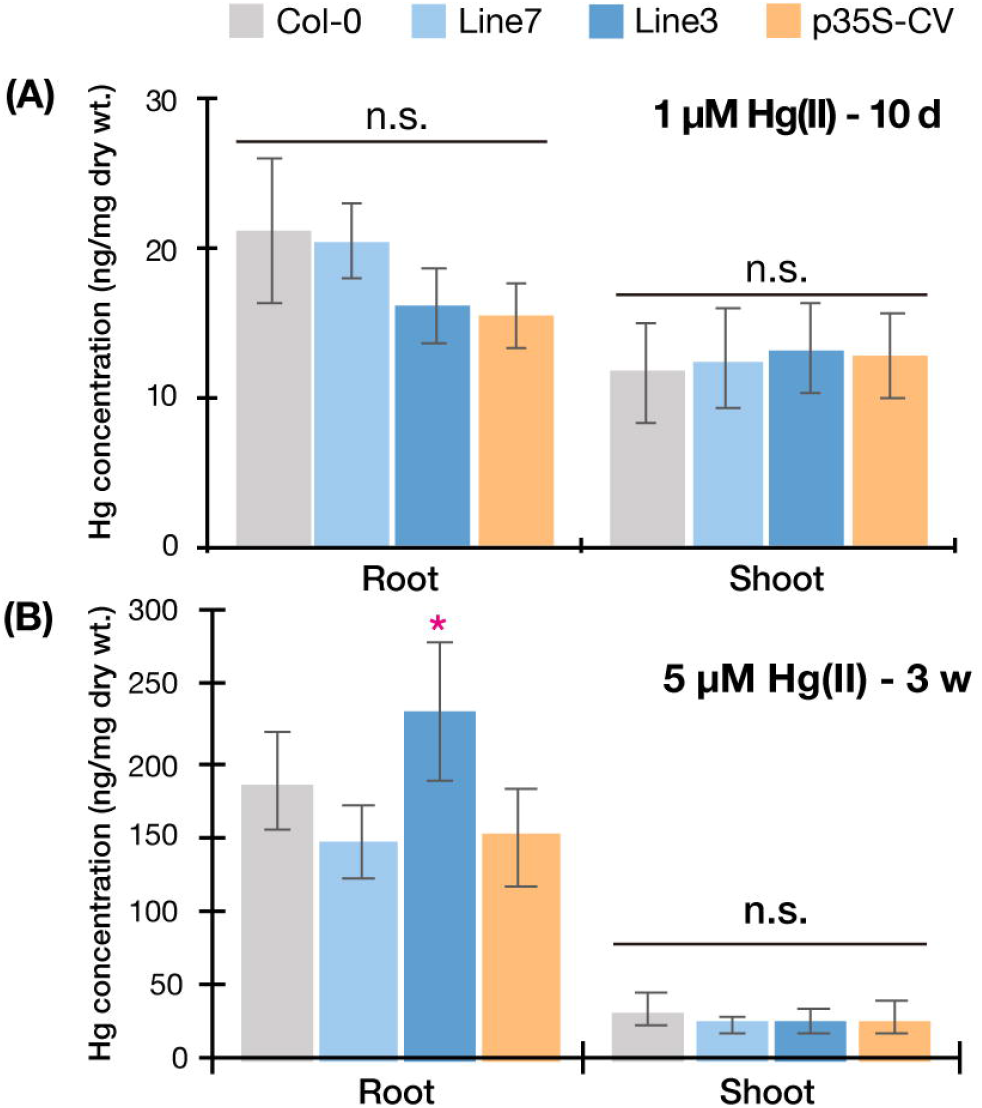
Mercury accumulation of Col-0, pRBCS1A-TCV lines, and p35S-CV line. Plants were exposed to 1 µM Hg(II) treatment for 10 d **(A)** and 5 µM Hg(II) treatment for 3 weeks **(B)**. Data represent means with SD from two independent experiments (*n* = 6 – 8). An asterisk indicates a significant difference from Col-0 (\**P* < 0.05, Dunnett’s test). n.s., not significant.

### Potentials of pRBCS1A-TCV lines for phytoremediation

A major significant finding of the mesophyll-specific MerC-AtVAM3 expression system is that MerC-AtVAM3 expression in shoots is sufficient for improving plant Hg(II) tolerance and that in roots is not essential. This point is significant for further molecular breeding towards phytoremediation which primarily requires higher metal accumulation in shoots. Vacuolar sequestration of toxic metals in the root cells can limit root-to-shoot translocation of toxic elements (Ueno et al., 2010; Miyadate et al., 2011; Uraguchi et al., 2018, 2020), which is not desirable for efficient phytoremediation. Although the ubiquitous expression of MerC-AtVAM3 did not inhibit shoot Hg accumulation under Hg(II) treatment (Figure 7), Hg was more retained in the roots of the p35S-CV line when exposed to methylmercury (Sone et al., 2017). Metal retention in roots derived from the active vacuolar sequestration may depend on the metal species and chemical forms, however, it is notable for phytoremediation that the shoot specific expression of MerC-AtVAM3 sufficiently enhances plant tolerance against at least Hg(II) stress without visible penalty of Hg accumulation in shoots.

We recently demonstrated that the root cell-type-specific expression of MerC-SYP121 enhanced Hg(II) uptake and lead to higher shoot Hg accumulation (Uraguchi et al., 2019a, 2019b). SYP121 is another Arabidopsis SNARE and plasma-membrane resident. MerC-SYP121 fusion protein is preferentially localized to the plasma-membrane and facilitates root Hg(II) uptake. Taking together with these three successful cases of the cell-type specific and organelle-targeted expression of MerC, pyramiding root-specific MerC-SYP121 expression and mesophyll-specific MerC-AtVAM3 expression systems would boost Hg uptake/accumulation ability in roots and Hg-detoxification activity in leaf mesophyll cells, respectively.

## Acknowledgments

We thank Prof. Dr. Angelika Mustroph (University of Bayreuth) for providing a plasmid containing the pRBCS1A fragment and the Arabidopsis Biological Resource Center (ABRC) for providing seeds of the p35S-GFP-δTIP line. This work was supported in part by the Japan Society for the Promotion of Science (Grant nos. 15H02839 and 18H03401 to M.K.).

## Author contributions

SU and MK designed the experiments. SU, YO, MO, SK, NY, and YuT conducted the experiments. SU, YO, RN, YaT, and MK analyzed the data. SU and MK wrote the manuscript, and all authors approved the manuscript.

