## Supplementary figures and images for "Mesophyll specific expression of a bacterial mercury transporter-based vacuolar sequestration machinery sufficiently enhances mercury tolerance of Arabidopsis"

### Supplementary Figure S1

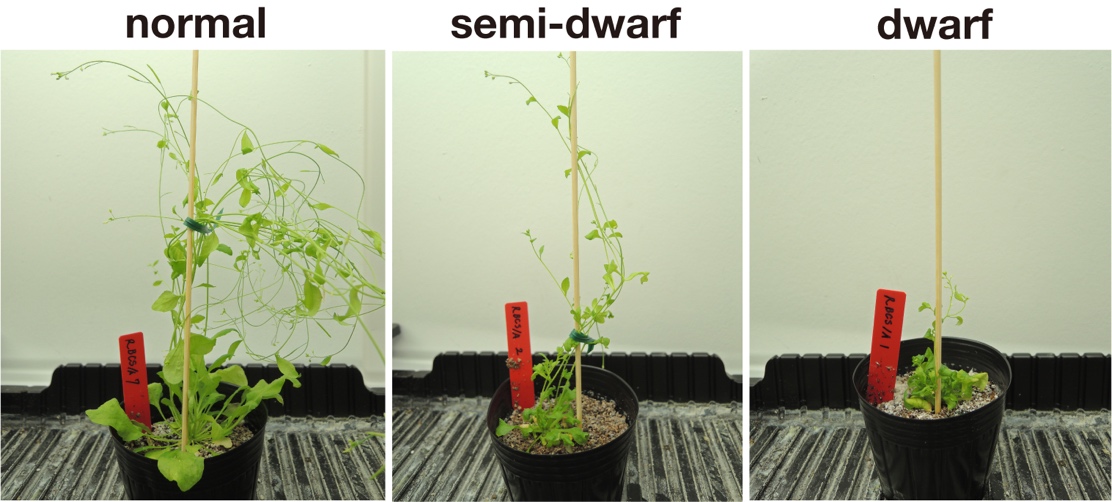

**Supplementary Figure S1.** Typical phenotypes of pRBCS1A-TCV transgenic T1 plants..
